# Defining the impact of rRNA processing on nucleolar organization and function

**DOI:** 10.1101/2025.10.19.682850

**Authors:** Maria Saraí Mendoza-Figueroa, Ellen Lavorando, Yulia Gonskikh, Charles Antony, Heidi Elashal, Anthony Yulin Chen, Hsin-Yao Tang, Dawn Carone, Vikram R. Paralkar, Kathy Fange Liu

## Abstract

The eukaryotic nucleolus is a highly organized, multilayered structure essential for ribosomal RNA (rRNA) processing and ribosome assembly. However, how the sequential steps of rRNA maturation, particularly the series of endonucleolytic cleavages, contribute to maintaining nucleolar architecture remains poorly understood. Here, we show that disruption of pre-rRNA processing, especially impaired cleavage of the 5′ external transcribed spacer (5′ETS), profoundly alters nucleolar organization. Specifically, defects in 5′ETS processing lead to the formation of a single large DAPI-negative nuclear structure and result in the mislocalization of nascent RNA, which diffuses throughout the disorganized nucleolus. These aberrant nucleoli exhibit a distinct proteomic profile, including downregulation of factors involved in splicing, cell cycle regulation, and chromatin organization, suggesting that the impact of nucleolar disorganization extends beyond ribosome biogenesis. Notably, we also observe mislocalization of heterochromatin markers, pointing to broader disruptions in nuclear architecture and gene regulation. Together, our findings reveal that proper 5′ETS cleavage is critical for preserving nucleolar compartmentalization and highlight the tight coupling between rRNA processing and nuclear organization.

## INTRODUCTION

Ribosome biogenesis is a fundamental cellular process for producing ribosomes, the molecular machines that synthesize proteins encoded by messenger RNA (mRNA)^1-4^. This complex process involves the coordination of numerous protein assembly factors and small nucleolar RNAs (snoRNAs) to process pre-ribosomal RNAs (rRNA) and assemble ribosome subunits within the nucleolus, a specialized subnuclear compartment^2,5,6^. The nucleolar sub-compartments are critical for stepwise ribosomal assembly and the outbound flux of maturing ribosomes^7,8^. The transcription and processing of rRNAs are essential for maintaining nucleolar structure. Inhibition of RNA Polymerase I disrupts rRNA synthesis, leading to significant rearrangements in nucleolar organization^9-11^. Similarly, disruption of rRNA processing or ribosome biogenesis influences nucleolar morphology^12-15^. However, the key steps and detailed mechanisms by which rRNA processing and ribosome assembly impact nucleolar organization and function are not understood.

The small (40*S*) and large (60*S*) subunits of the ribosome are assembled by their dedicated processomes (ribonucleoprotein complexes that serve as platforms for their assembly and maturation^16,17^). Processomes provide a structured environment for the ordered folding and modification of rRNA and for the incorporation of ribosomal proteins to form functional and competent ribosomes for protein synthesis^16,17^. The small subunit (SSU) processome forms around the nascent pre-rRNA molecule in the early stages of small subunit assembly^7^. The human SSU processome comprises over 70 protein factors and several snoRNAs, including U3 snoRNP, which plays a crucial role in rRNA processing^7^. The rRNA precursor undergoes a series of cleavage events that remove 5’ and 3’ external transcribed spacers (5’ETS and 3’ETS) and internal transcribed spacers 1 and 2 (ITS1 and ITS2) from the pre-rRNA, generating the mature 18*S* rRNA incorporated into the 40*S* subunit^7^. The human 5’ETS notably has a length of 3.6 kilobases, but only 25% of the total sequence is necessary to produce mature 18*S* rRNA when using a synthetic nucleotide template^7^. The function of the remaining 75% of the 5’ETS remains unclear. The presence or absence of the full-length 5’ETS distinguishes the three maturation states of the human SSU processome: pre-A1, pre-A1*, and post-A1 stages. States pre-A1 and pre-A1* are characterized by the presence of the 5’ETS, whereas the post-A1 state contains only a minimal ordered segment of the 5’ETS near the interacting UtpA and UtpB complexes^16,18^. The large subunit (LSU) processome emerges later in the assembly pathway, surrounding the maturing 28*S* pre-rRNA molecule. The pre-rRNA undergoes cleavage and modification, ultimately generating the mature 28*S* rRNA and 5.8*S* rRNA. Together with 5*S* rRNA assembled by 5S ribonucleoprotein, they form the functional 60*S* subunit^19^. Disruptions in either SSU or LSU processome function can lead to ribosome deficiency and protein synthesis defects, which often underlie diseases (referred to as ribosomopathies) with varying degrees of severity^20-22^.

Most steps of ribosome biogenesis occur in the three highly organized sub-compartments of the nucleolus^5^. These three layers are fibrillar centers (FCs), dense fibrillar components (DFCs), and granular components (GCs)^5^. The FCs are enriched with RNA polymerase I subunits responsible for transcribing rRNA. Transcription of rRNA starts mainly at the borders between the FCs and DFCs. After being transcribed, pre-rRNAs are further processed in the DFCs before moving to the GC layer for final maturation (**Fig. 1a**). Previous studies suggest that the formation of the distinct layers of the nucleolus is driven by a biophysical process named phase separation^23,24^. This process involves a complex network of interactions between rRNA molecules and the proteins of the nucleolus. Importantly, the nucleolus is the largest and most prominent nuclear body. It forms around ribosomal DNA (rDNA) clusters located on the short arms of the five human acrocentric chromosomes 13, 14, 15, 21, and 22^25^. Beyond its role in ribosome biogenesis, the nucleolus anchors regions of constitutive and facultative heterochromatin, including those marked by H3K9me3 and H3K27me3^26^. Given this structural and functional integration, changes in nucleolar organization may have broad consequences for nuclear architecture, including alterations in heterochromatin distribution and gene regulation.

**Figure 1.**
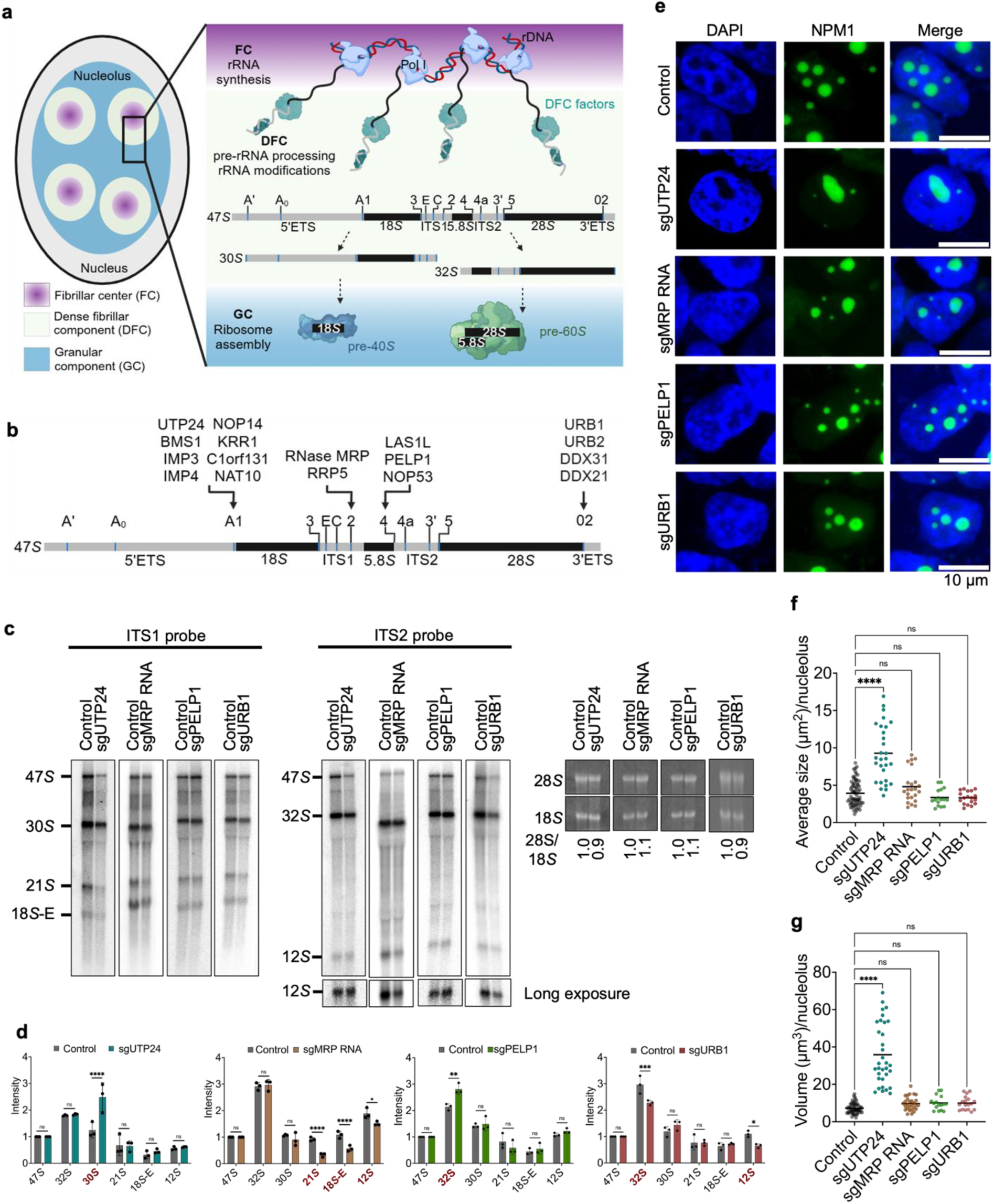
Specific pre-rRNA processing factors are critical for nucleolar morphology. (**a**) Schematic of the mammalian nucleolus illustrating its three sub compartments: the fibrillar center (FC), dense fibrillar component (DFC), and granular component (GC). rRNA transcription initiates at the FC/DFC boundary, followed by pre-rRNA processing in the DFC and ribosomal subunit assembly in the GC. Key molecular components and major steps in ribosome biogenesis are depicted. (**b**) Diagram of the 47*S* pre-rRNA transcript showing cleavage sites and associated processing steps. Representative protein and RNA factors involved at each stage of pre-rRNA processing are indicated. (**c**) Representative Northern blot analyses of pre-rRNA processing in HEK293T cells depleted of the indicated ribosome assembly factors using CRISPR-Cas9 with paired sgRNAs. Probes targeting the ITS1 and ITS2 regions were used to detect processing intermediates. Mature 28*S* and 18*S* rRNAs were visualized by methylene blue staining. Ratios of 28*S*/18S rRNA are indicated. Membranes were overexposed to enhance detection of the 12*S* intermediate. Each experiment was performed in triplicate. (**d**) Quantification of rRNA processing intermediates from panel (c), normalized to the 47*S* precursor. Data are from three independent biological replicates. Intermediates associated with processing defects are highlighted in bold red. Statistical analysis was performed using two-way ANOVA; ****p < 0.0001, ***p = 0.0002, **p = 0.0014, *p = 0.0164, ns = not significant (α > 0.05). Mean ± SD are shown. (**e**) Representative immunofluorescence images of control and CRISPR-Cas9–edited HEK293T cells depleted of UTP24, MRP RNA, PELP1, or URB1. Nuclei are stained with DAPI (blue); nucleoli are labeled with anti-NPM1 (GC marker; green). Scale bar, 10 µm. (**f**) Quantification of nucleolar area (µm²) from panel (e). Data were analyzed using the Kruskal–Wallis test; ****p < 0.0001, ns = not significant (α > 0.05). Mean values are shown. (**g**) Quantification of nucleolar volume (µm³) from panel (e), analyzed as in panel (f). ****p < 0.0001, ns = not significant (α > 0.05). Mean values are shown.

Here, we systematically examine how key pre-rRNA processing steps contribute to nucleolar organization and their impacts on heterochromatin marker distribution. We find that depletion of ribosome biogenesis factors involved in 5′ETS cleavage in human cells leads to profound disruption of nucleolar architecture. Proximity-labeling proteomics reveals that disorganized nucleoli exhibit distinct RNA and protein compositions compared to functional nucleoli. These structural alterations are accompanied by a redistribution of the heterochromatin mark H3K27me3. Together, our findings demonstrate that 5′ETS processing is critical for proper nucleolar assembly and highlight how defects in rRNA maturation can reshape both the nucleolar interactome and nuclear chromatin organization.

## RESULTS

### Depletion of 5′ETS cleavage factors more robustly induces large, DAPI-negative structures compared to depletion of ITS1/2 or 3′ETS cleavage factors

The 47*S* pre-rRNA is processed through a series of cleavage events to generate precursors of the mature 18*S*, 5.8*S*, and 28*S* rRNAs (**Fig. 1a**). To examine how disruption of specific processing steps affects nucleolar morphology, we used a CRISPR-Cas9–based strategy to deplete individual proteins or RNA factors involved in distinct stages of rRNA maturation^27-31^. Specifically, we targeted: (1) UTP24, which mediates 5′ETS cleavage^32-34^; (2) MRP RNA, which facilitates ITS1 site 2 cleavage to separate precursors destined for the small and large subunits^28,35^; (3) PELP1, which promotes ITS2 cleavage [28]; and (4) URB1, which is required for 3′ETS processing^31^ (**Fig. 1a, 1b**). Genomic PCR confirmed high depletion efficiency (80–92%) for each target (**Extended Data Fig. 1a**).

To validate rRNA processing defects, we performed Northern blot analyses in biological triplicates using probes targeting ITS1 and ITS2 regions (**Fig. 1c; Extended Data Fig. 1b**). As expected, UTP24 depletion led to accumulation of the 30*S* intermediate (ITS1 probe), while MRP RNA depletion caused a reduction in 21*S*, 18*S*-E, and 12*S* rRNA intermediates (detected with both ITS1 and ITS2 probes). PELP1 depletion resulted in accumulation of 32*S* rRNA, and URB1 depletion reduced levels of both 32*S* and 12*S* rRNA intermediates (ITS2 probe) (**Fig. 1c, 1d**). These results are consistent with previous studies that successful depletion of annotated rRNA-processing factor led to specific defects in rRNA processing steps^28,29,31-35^.

We next assessed how these disruptions affect nucleolar morphology using immunofluorescence. Depletion of UTP24 led to the formation of a prominent single, enlarged DAPI-negative region marked by nucleophosmin (NPM1) (**Fig. 1e**). Quantification revealed that nucleolar area increased approximately 2.4-fold in UTP24-depleted cells compared to controls (control mean: 3.9 µm²; sgUTP24: 9.3 µm²) (**Fig. 1f**), with nucleolar volume increasing ∼4.9-fold (control: 7.2; mutant: 35.8) compared to control cells (**Fig. 1g**). In contrast, depletion of MRP RNA, PELP1, or URB1 did not result in significant changes in nucleolar size (**Fig. 1e–1g**).

To further assess this specificity, we depleted additional ITS1, ITS2, and 3′ETS processing factors -RRP5^36^, NOP53^30^, URB2, DDX21, and DDX31^31^- using the same strategy (**Extended Data Fig. 1c, 1d**). None of these depletions significantly altered nucleolar area or volume (**Extended Data Fig. 1e, 1f**). Together, these results indicate that disruption of 5′ETS processing, compared to other rRNA processing steps, strongly impacts nucleolar size and volume, suggesting a unique role for this early processing event in shaping nucleolar architecture.

### Depletion of protein assembly factors involved in 5’ETS processing severely disrupts nucleolar organization

We next asked whether the single large DAPI-negative region observed upon UTP24 depletion is specific to UTP24 or reflects a general consequence of impaired 5’ETS cleavage. To identify additional ribosome assembly factors involved in A1 site cleavage, we examined components of the human small subunit (SSU) processome^16^. In addition to UTP24, we depleted other proteins proposed to directly mediate A1 site cleavage within the pre-A1 SSU complex - BMS1, IMP3, IMP4, NOP14, KRR1, C1orf131^16^, and NAT10^37^ (**Extended Data Fig. 2a**). Efficient depletion of each factor (**Extended Data Fig. 2b, 2c**) led to an accumulation of the 30*S* rRNA intermediate (**Fig. 2a, 2b; Extended Data Fig. 2d**), consistent with defective A1 cleavage. Notably, depletion of any of these proteins also resulted in a prominent single, enlarged DAPI-negative region (**Extended Data Fig. 2e**), phenocopying the effect of UTP24 depletion and reinforcing a link between A1 cleavage and nuclear morphology. Despite belonging to distinct SSU subcomplexes^38^, all factors examined yielded a consistent morphological phenotype.

**Figure 2.**
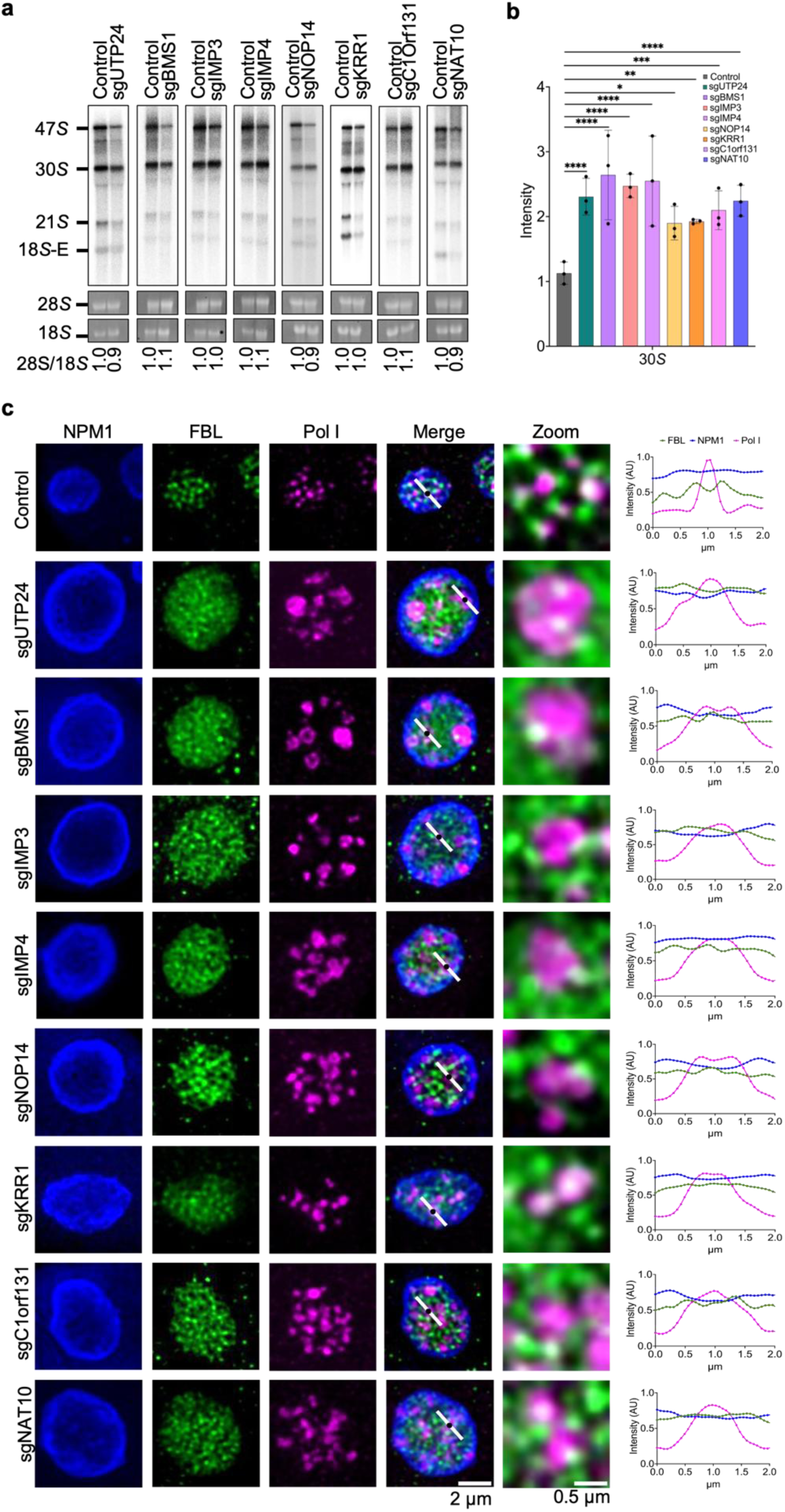
Pre-rRNA processing factors involved in A1 site cleavage are essential for nucleolar organization. (**a**) Representative Northern blots showing the effects of depleting ribosomal processing factors involved in A1 site cleavage. Depletion was achieved via CRISPR-Cas9 using paired sgRNAs. Loss of individual factors consistently resulted in the accumulation of the 30*S* intermediate, detected using a DNA probe targeting the ITS1 region. Mature 28*S* and 18*S* rRNAs were visualized by methylene blue staining; 28*S*/18*S* ratios are indicated. Each blot represents one of three independent biological replicates. (**b**) Quantification of the 30*S* rRNA intermediate from panel (a). Signal intensities were measured across three independent replicates using Fiji, and values were normalized to the corresponding 47*S* precursor. Statistical significance was determined by two-way ANOVA; ****p < 0.0001; ***p = 0.0004; **p = 0.0076; *p = 0.0113; ns (not significant) > 0.05. Data represent mean ± standard deviation. (**c**) Immunofluorescence analysis by super-resolution microscopy of nucleolar organization in HEK293T cells transfected with pX459 control vector or pX459 containing sgUTP24 targeting the indicated A1 cleavage factors. Cells were stained for markers of the nucleolar subcompartments: fibrillar center (FC, α-RPA194/Pol I), dense fibrillar component (DFC, α-fibrillarin/FBL), and granular component (GC, α-NPM1). Fluorescence intensity profiles were generated by plotting signal distributions centered on Pol I centroids (white line in merged panels; Pol I centroid indicated by black dot). Average fluorescence intensities (arbitrary units, AU) are shown at right. Scale bars: 2 µm (merged panels), 0.5 µm (zoomed panels).

To investigate how impaired A1 cleavage impacts nucleolar sub-compartmentalization, we assessed the multilayered organization of the nucleolus by staining for classical compartment markers: RPA194 (Pol I) for the fibrillar center (FC), fibrillarin (FBL) for the dense fibrillar component (DFC), and NPM1 for the granular component (GC) (**Fig. 2c**)^39^. Using super-resolution microscopy, we quantified the spatial distribution of these markers relative to Pol I signal centroids (white line, merged panels, **Fig. 2c**). In control cells, Pol I clusters were surrounded by FBL and NPM1, consistent with the classical nucleolar architecture previously described^40^. In contrast, depletion of any of the A1 cleavage factors led to remarkable disorganization of nucleolar layers. Specifically, fluorescence intensity profiles revealed that both Pol I and FBL signals were significantly broadened, indicating that the FC and DFC layers are more dispersed in the enlarged nucleoli (**Fig. 2c**). These results highlight a critical role for ribosome assembly factors involved in A1 site cleavage in maintaining the structural integrity of the FC and DFC layers within the nucleolus.

To determine whether the observed phenotype stems specifically from disrupted A1 site cleavage, as opposed to indirect effects of SSU complex perturbation, we used a complementary strategy. We expressed a catalytically inactive mutant of UTP24, which normally cleaves the 5′ETS at sites A0, A1, and E in human cells (**Fig. 3a**)^34^. HEK293T cells were transduced with lentivirus encoding HA-tagged D72N/D142N UTP24 or an empty vector control, achieving high transduction efficiency (**Extended Data Fig. 3a, 3b**). Western blotting confirmed expression of the mutant protein (**Extended Data Fig. 3c**), and Northern blot analysis revealed a robust accumulation of the 26*S* rRNA precursor (**Fig. 3b–3d**), consistent with previous reports^34^. Expression of catalytically inactive HA-D72N/D142N UTP24 induced a clear nucleolar enlargement, with an average nucleolar area increase of ∼2.3-fold compared to control cells (control: 4.26 µm²; mutant: 9.81 µm²) (**Fig. 3e**), and an average volume increase of ∼ 4.8-fold (control: 7.96; mutant: 38.56) (**Fig. 3f**), changes comparable to those seen with UTP24 depletion (**Fig. 1e, 1f**). Additionally, we observed an expansion of the FC and DFC layers in these enlarged nucleoli, phenocopying the effects of UTP24 and other 5′ETS processing factor depletions (**Figs. 2c, 3e, 3g**). Together, these results support a model in which proper removal of the 5′ETS during early pre-rRNA processing is essential for maintaining nucleolar organization.

**Figure 3.**
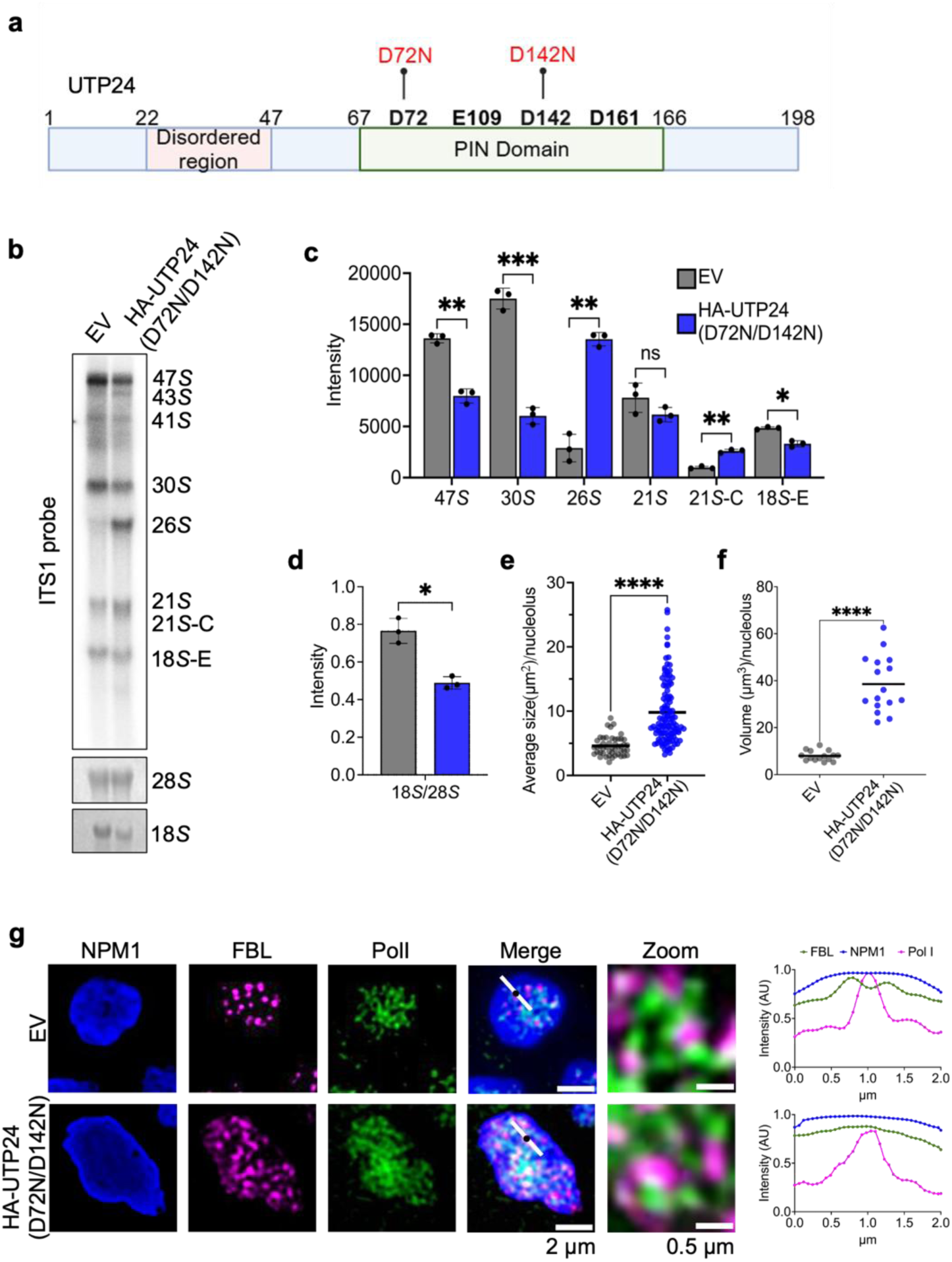
The endonuclease activity of UTP24 is essential for nucleolar organization. (**a**) Domain architecture of human UTP24, indicating the disordered region, PIN domain, and catalytic residues. Point mutations D72N and D142N were introduced to generate a catalytically inactive form of UTP24. (**b**) Northern blot analysis showing that expression of catalytically inactive HA-tagged UTP24(D72N/D142N), introduced via lentiviral transduction, results in accumulation of 26*S* and 21*S*-C pre-rRNA intermediates. Detection was performed using a probe against the ITS1 region. EV, empty vector. (**c**) Quantification of rRNA precursor bands from three independent Northern blot replicates, analyzed using Fiji. Mature 28*S* and 18*S* rRNAs were visualized by methylene blue staining. Statistical analysis was performed using two-way ANOVA; ***p < 0.001; **p < 0.01; *p < 0.1; ns = not significant (α > 0.05). Data represent mean ± standard deviation. (**d**) Quantification of 28*S* and 18*S* rRNA band intensities from panel (b). Expression of catalytically inactive UTP24(D72N/D142N) specifically reduces 18*S* rRNA levels. The data represent three biological replicates. Paired t-test; *p < 0.01; ns = not significant (α > 0.05). Mean ± standard deviation is shown. (**e**) Quantification of nucleolar area (µm²) from panel (f). Statistical analysis was performed using an unpaired two-tailed t-test; ****p < 0.0001. Mean values are shown. (**f**) Quantification of nucleolar volume (µm³) from panel (g). Statistical analysis was performed using an unpaired two-tailed t-test;****p < 0.0001 (α > 0.05). Mean values are shown. (**g**) Immunofluorescence analysis by super-resolution microscopy of nucleolar substructures in HEK293T cells transduced with either empty vector (EV) or HA-UTP24(D72N/D142N). Cells were stained with antibodies against the fibrillar center (FC, α-RPA194/Pol I), dense fibrillar component (DFC, α-FBL), and granular component (GC, α-NPM1). Fluorescence intensity profiles were generated by plotting marker distributions centered on Pol I signal centroids (white line in merged panels; Pol I centroid indicated by a black dot). Average fluorescence intensities (arbitrary units, AU) are shown at right. Scale bars: 2 µm (merged panels), 0.5 µm (zoomed panels).

### Defective 5′ETS cleavage disrupts nucleolar dynamics

Nucleolar components are highly dynamic, and interactions between nucleolar proteins and RNA undergo phase separation, resulting in non-intermixed, liquid-like phases^41^. This immiscibility has been proposed to drive the formation of the multilayered nucleolar architecture^5,24^. As shown above, upon UTP24 depletion, the dense fibrillar component and the fibrillar center layers of the enlarged nucleoli are redistributed (**Fig. 2**). We next asked whether nucleolar dynamics are also impaired in these enlarged nucleoli. To address this, we analyzed the dynamics of the granular component protein NPM1 using fluorescence recovery after photobleaching (FRAP). An mCherry-tagged NPM1 was expressed in HEK293T cells depleted of UTP24 (sgUTP24) and compared to the knockdown control cells. FRAP signals were then recorded (**Fig. 4**). In control cells, mCherry-NPM1 exhibited rapid recovery, reaching maximal fluorescence intensity approximately 5.3 seconds after bleaching. In contrast, in UTP24-depleted cells, recovery was delayed, reaching maximum intensity at ∼8.6 seconds post-bleach (**Fig. 4a, 4b**). The mobile fraction of mCherry-NPM1 was markedly reduced in UTP24-depleted cells (Control: 0.8; sgUTP24: 0.2), and the half-time of recovery (t_1/2_) was 1.6-fold longer (Control = 2.6 s; sgUTP24: 4.3 s). These findings indicate that nucleolar dynamics are significantly slower under UTP24 depletion, suggesting that the liquid-like properties of nucleoli are compromised.

**Figure 4.**
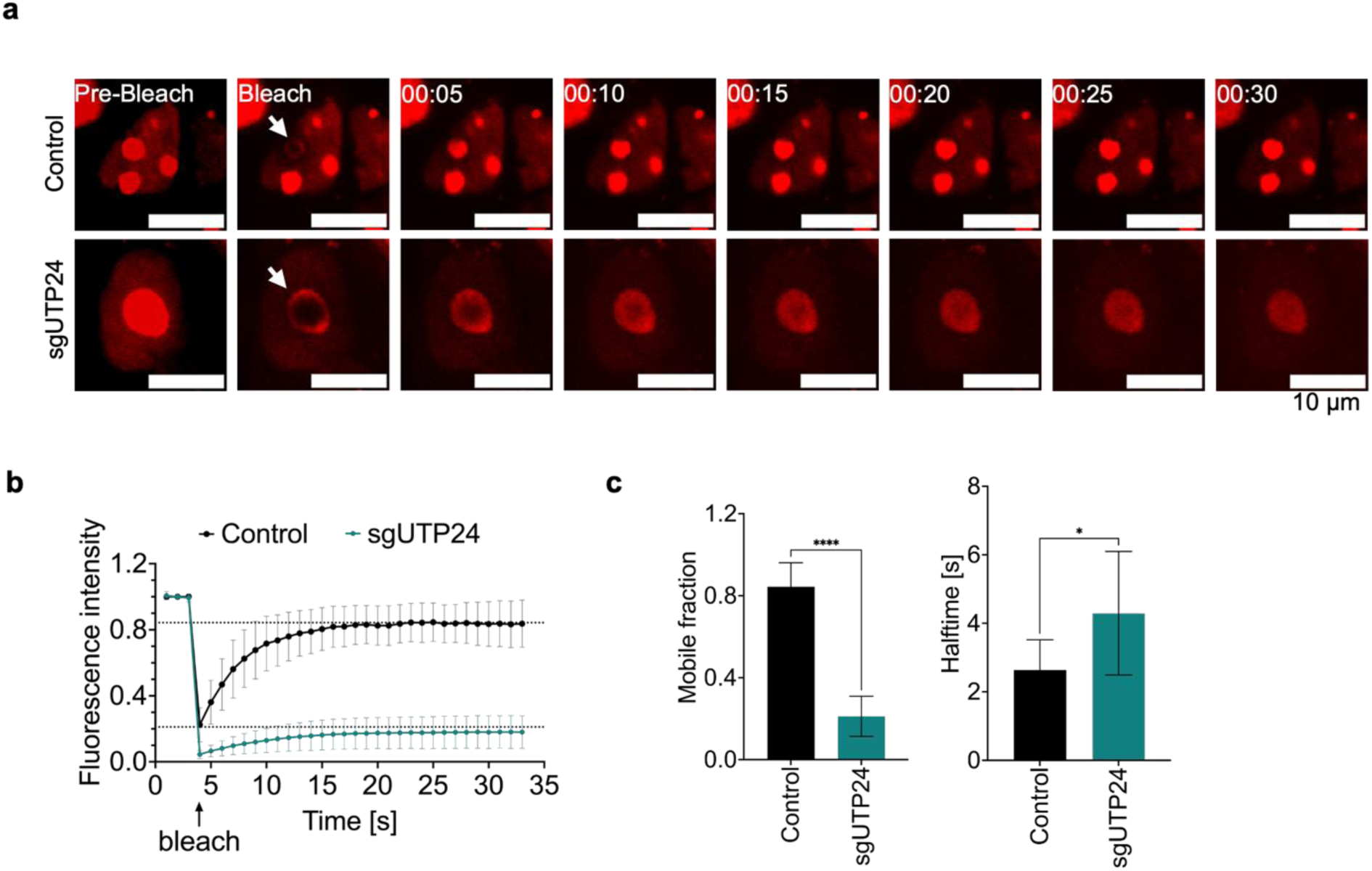
Depletion of UTP24 impairs nucleolar dynamics. (**a**) Fluorescence recovery after photobleaching (FRAP) assays were performed in HEK293T cells co-transfected with either pX459 control vector or pX459 containing sgUTP24, along with mCherry-tagged NPM1. A defined nucleolar region was photobleached (arrows), and fluorescence recovery was recorded over time (in seconds). Scale bar: 10 µm. (**b**) Quantification of mCherry-NPM1 fluorescence intensity over time, corresponding to panel (a). The bleach time point is indicated. Horizontal dotted lines represent the mobile fraction (fluorescence plateau after recovery). (**c**) Quantification of mobile fraction (left) and recovery half-time (t₁/₂, right) reveals significantly slower recovery kinetics of mCherry-NPM1 in UTP24-depleted cells. Statistical analysis was performed using one-way ANOVA: ****p < 0.0001; *p < 0.05; ns = not significant (α > 0.05). Data represent mean ± standard deviation.

### Defects in 5’ETS cleavage disrupt RNA distribution and nucleolar interactome

Since nucleolar organization is critical for the processing and flux of newly synthesized rRNA^7^, we assessed the impact of depletion of UTP24 on nascent RNA synthesis using 5-ethynyluridine (5-EU) incorporation followed by click chemistry in HEK293T cells^7,42,43^ (**Fig. 5a**). In control cells, most of the 5-EU-labeled RNA signals colocalized with RNA polymerase I (Pol I) within nucleoli, consistent with previous observations^7,42,43^. Upon UTP24 deletion, however, nascent RNA exhibited a diffuse and expanded distribution throughout the disorganized nucleoli (**Fig. 5a**). Quantification of total Pol I signal intensity in morphologically normal nucleoli of control cells versus the enlarged nucleoli in UTP24-depleted cells revealed no significant differences (**Extended Data Fig. 4a**), suggesting unaltered Pol I protein levels upon UTP24 depletion, which is consistent with western blot quantifications (**Fig. 5e, 5f**). Additionally, the total nuclear 5-EU signal decreased (**Fig. 5b**), suggesting that transcriptional output is reduced, likely due to a slower rate of rRNA synthesis. These results, together with the reduced levels of 47*S* pre-rRNA precursors (**Extended Data Fig. 4b**), indicate that rRNA transcription is attenuated despite unchanged Pol I abundance. Nonetheless, we cannot exclude the possibility that the observed reduction in global nascent RNA arises from impaired transcription by other RNA polymerases.

**Figure 5.**
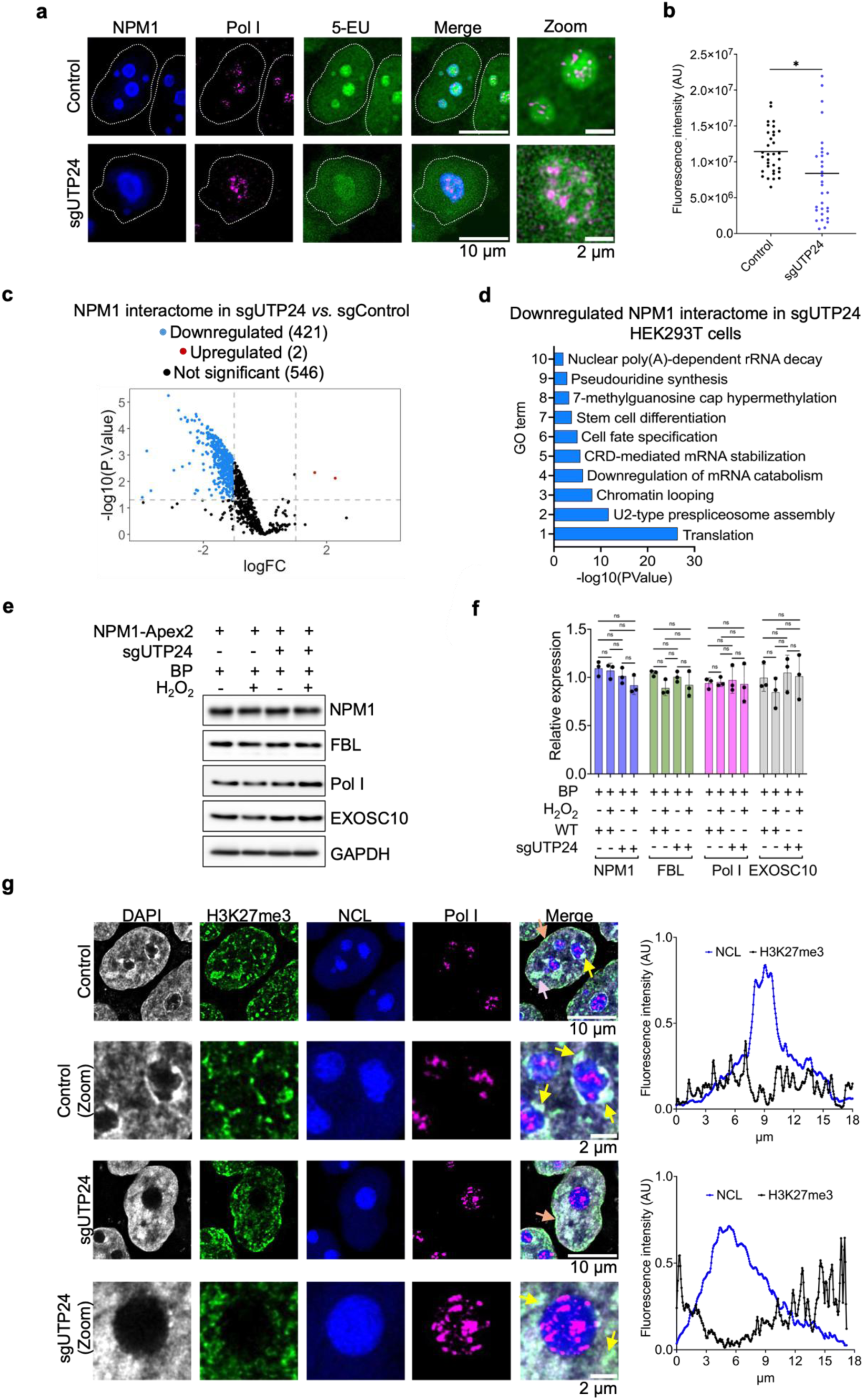
UTP24 depletion disrupts nucleolar protein interactome and heterochromatin organization. (**a**) Immunofluorescence analysis by super-resolution microscopy of nascent RNA labeled with 5-ethynyl uridine (5-EU) in HEK293T cells transfected with either pX459 control vector or pX459 containing sgUTP24. Nucleolar substructures were visualized using antibodies against the fibrillar center (FC; α-RPA194/Pol I) and the granular component (GC; α-NPM1). Scale bars: 2 µm (merged panels), 0.5 µm (zoomed panels). (**b**) Quantification of nuclear 5-EU fluorescence intensity from panel (a). Unpaired t-test; ns = not significant. Mean values are shown. (**c**) Volcano plot of differentially biotinylated proteins identified by streptavidin pull-down following proximity labeling with Nucleophosmin-Apex2 in sgUTP24 versus cells transfected with pX459 control vector. Proteins with *p* < 0.05 and log₂ fold change (logFC) ≥ 1 are shown in red; ≤ –1 in blue; non-significant proteins in black. (**d**) Gene Ontology (GO) enrichment analysis of significantly downregulated proteins (blue dots from panel (c) in the NPM1 interactome upon UTP24 depletion. (**e**) Western blot analysis of nucleolar proteins (NPM1, FBL, and Pol I) and the RNA surveillance factor EXOSC10 in lysates from control and sgUTP24 cells overexpressing Nucleophosmin-Apex2. GAPDH was used as a loading control. (**f**) Quantification of protein levels shown in panel (e). One-way ANOVA; ns = not significant (α > 0.05). Data represent mean ± standard deviation. (**g**) Immunofluorescence analysis of the heterochromatin marker H3K27me3 (α-H3K27me3), FC (α-RPA194/Pol I), and GC (α-Nucleolin/NCL) in control and sgUTP24 cells. Scale bar: 10 µm. Zoomed panels highlight representative H3K27me3 distribution around the nucleolus; scale bar: 2 µm. Quantification of H3K27me3 and NCL fluorescence intensities is shown at right. Enrichment of H3K27me3 at distinct nuclear loci is indicated: nucleolus (yellow arrows), nuclear periphery (orange arrows), and regions not associated with either compartment (pink arrows).

To explore the impact of nucleolar disorganization upon impaired 5’ETS cleavage on the proteomic landscape, we employed APEX2-based proximity labeling using an NPM1-APEX2 construct^44,45^ (**Extended Data Fig. 4c**). After validating UTP24 depletion (**Extended Data Fig. 4d**), we compared the NPM1 interactome in wild-type versus UTP24-depleted cells. Streptavidin-HRP immunoblots confirmed efficient biotinylation and pulldown of proximal proteins (**Extended Data Fig. 4e**). Mass spectrometry analysis revealed a reduction of 421 proteins in the NPM1 interactome following UTP24 depletion (**Fig. 5c; Extended Data Fig. 4f, Table S2**), based on detection by at least two peptides in a minimum of two biological replicates. Gene Ontology analysis showed that downregulated proteins participate in diverse pathways, including translation (ribosomal proteins), U2-type spliceosome assembly (SF3A/SF3B complexes), chromatin remodeling and looping (RUVBL1–RUVBL2, DEAD/DEAH-box helicases), and RNA turnover (nuclear exosome) (**Fig. 5c, 5d**). Interestingly, while proteins such as the exosome component EXOSC10 were reduced in the nucleolar interactome, their total cellular levels remained unchanged, similar as observed in other nucleolar markers, including Pol I (FC), fibrillarin (DFC), and NPM1 (GC) (**Fig. 5e, 5f**). These findings suggest that impaired 5’ETS cleavage leads to nucleolar disorganization, which broadly alters the composition and function of the nucleolar proteome beyond canonical roles in ribosome biogenesis.

### The heterochromatin marker H3K27me3 redistributes upon impaired 5′ETS cleavage

The nucleolus, the largest membraneless nuclear body, serves as an anchor for heterochromatin regions. These regions, termed nucleolus-associated domains (NADs), are distributed across all chromosomes and are primarily located near telomeres, centromeres, and the nucleolar organizer regions (NORs) of the five acrocentric chromosomes (13, 14, 15, 21, and 22)^46^, Thus the nucleolus contributes to the three-dimensional organization of genome architecture. Given the dramatic changes in nucleolar organization upon impaired 5′ETS cleavage, we investigated how these structural alterations affect heterochromatin organization by examining the distribution of the repressive heterochromatin mark H3K27me3 using immunofluorescence (**Fig. 5g; Extended data Fig. 4g**). In control cells, H3K27me3 localizes to the nucleolar periphery (yellow arrow, **Fig. 5g**) and is also enriched at the nuclear periphery (orange arrow, **Fig. 5g**), consistent with prior findings^46-48^. Additionally, it accumulates in other nuclear regions not associated with either location (pink arrows, **Fig. 5g**). Upon UTP24 depletion, H3K27me3 localization at the nuclear periphery remained largely unchanged (orange arrows, **Fig. 5g**). However, around the enlarged nucleoli, the signal appeared more diffuse, losing the sharply defined distribution seen in controls (zoom-in panels, **Fig. 5g**). These changes were also evident in fluorescence intensity plots (**Fig. 5g; Extended Data Fig. 4g**). Together, these results suggest that disruption of 5′ETS processing alters nucleolar organization and, consequently, redistributes nucleolus-associated heterochromatin, potentially affecting gene regulation.

## DISCUSSION

Nucleolar organization and dynamics are critical for ribosome biogenesis^5,7,40^. Our study delves into the intricate relationship between rRNA processing and nucleolar organization, building upon the established role of the nucleolus in facilitating ribosome assembly. While previous studies have highlighted the nucleolus’s crucial role in ribosome biogenesis and flux^5,7,40^, the current work illustrates that rRNA processing, specifically^34^ an early ribosome biogenesis step, 5’ETS removal, impacts the nucleolus’s structural integrity, actively shapes the organization of this subnuclear compartment, and further influencing heterochromatin marker distribution (**Fig. 6**). This work is protein-centric, focusing on the ribosome assembly factors involved in key steps of rRNA processing, to further our understanding of how rRNA processing influences nucleolar organization and function. Together with previous studies^5,7,31,40^, we demonstrate the dependence of the multi-layered nucleolar architecture on the fidelity of rRNA processing.

**Figure 6.**
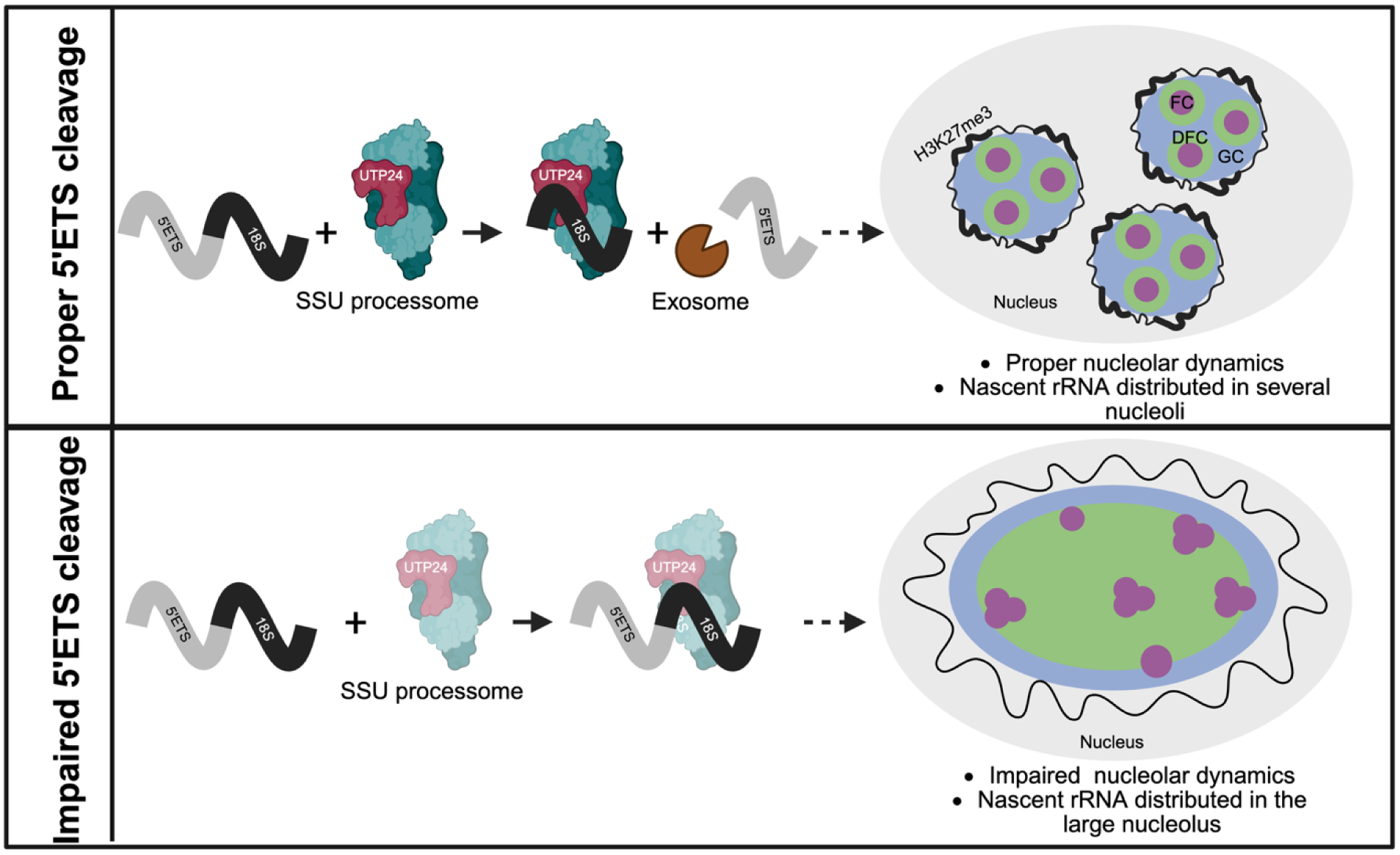
5′ETS cleavage by the SSU processome shapes nucleolar and chromatin architecture. **Top:** During ribosome biogenesis, the SSU processome cleaves the 5′ external transcribed spacer (5′ETS) at the A1 site via the endonuclease UTP24. The released 5′ETS is then degraded by the RNA exosome. Efficient 5′ETS removal enables proper rRNA processing and promotes nucleolar organization, with clear separation of the fibrillar center (FC), dense fibrillar component (DFC), and granular component (GC). Proper nucleolar organization in turn supports the establishment of perinucleolar heterochromatin domains. **Bottom:** In the absence of efficient 5′ETS cleavage, the 30S pre-rRNA intermediate accumulates, leading to nucleolar enlargement and disrupted nucleolar architecture. FCs form enlarged aggregates, and DFC components become mislocalized throughout the nucleolus. These structural alterations in the nucleolus are accompanied by global changes in heterochromatin distribution, affecting nuclear chromatin organization.

Among the 200 ribosome assembly factors and the complex pre-rRNA processing steps, we demonstrate that depleting proteins involved in 5’ETS cleavage consistently led to disorganized nucleoli, underscoring the critical role of 5’ETS removal. However, depletion of these assembly factors also perturbs the protein scaffold of ribosomal sub-complexes associated with specific rRNA regions: 5’ETS, 5’, central, 3’ major, and 3’ minor domains. Therefore, an alternative explanation is that depleting such proteins hinders proper SSU formation rather than the 5’ETS cleavage itself and disrupts nucleolar organization. To specifically impair 5′ETS cleavage while preserving the SSU protein scaffold, we expressed a catalytically inactive UTP24 mutant^34^. The results recapitulated the nucleolar disorganization observed upon depletion of other factors involved in 5′ETS cleavage (**Fig. 3**). While we cannot fully exclude the possibility that re-expression of the mutant UTP24 alters SSU assembly than wild-type UTP24 expression, our findings support that rRNA cleavage at the 5′ETS has a strong impact on proper nucleolar organization.

Two recent studies support our finding on the importance of 5’ETS cleavage on nucleolar organization^49,50^. Pan *et al.*^47^, using antisense oligonucleotides to block 5’ETS processing, demonstrated that inhibiting cleavage at either the A0 or A1 sites leads to expanded FC/DFC regions. Quinodoz *et al*.^46^, employing engineered synthetic nucleoli, showed that blocking U3 snoRNA-mediated 5’ETS cleavage led to FC/DFC regions displaced outside the GC layer. These two complementary studies reinforce the importance of 5’ETS cleavage in maintaining nucleolar architecture and provide high-resolution imaging of rRNA intermediates within the three nucleolar sub-compartments. Although our work also supports the conclusion that 5’ETS cleavage led to nucleoli organization, we do not observe FC/DFC layer redistributing to outside of the GC layers^49^. Our work adds to these findings by emphasizing the critical role of SSU protein assembly factors in rRNA processing and nucleolar organization, offering a protein-centric perspective that complements these exciting advancements. These three studies consistently support that the pre-rRNA with unprocessed 5’ETS generates the large DAPI-negative nuclear structures, which also suggests a link between early ribosome biogenesis steps and nucleolar organization, introducing complexity to the regulatory network governing nucleolar dynamics.

Another important finding our study provides is mapping the protein interactome within the large DAPI-negative nuclear structures when 5’ETS is not properly processed. Dysregulation induced by disrupted 5’ETS cleavage induces profound changes in the protein network of the nucleus. Our APEX2-MS proximity proteomics data show significant alterations in DNA repair, chromatin looping, and cell cycle regulation pathways, suggesting implications for genomic stability and cellular homeostasis. Future research should unravel the mechanistic insights into how dysregulated rRNA processing goes beyond influencing translation to induces changes in nuclear protein networks to provide a more comprehensive understanding of the functions of rRNA processing in gene regulation. Further investigations are also needed to explore the functional consequences of dysregulation in DNA repair and cell cycle regulation pathways, shedding light on the broader impact on cellular function and potential implications for disease states associated with aberrant ribosome biogenesis.

Lastly, our work shows that depleting proteins involved in ITS2 and 3’ETS cleavages (in LSU biogenesis) also affect the nucleolar organization, albeit to a lesser extent than 5’ETS retention (**Fig. 1e and Extended Data Fig. 1d**). A recent study using NPM1 as a nucleolar marker and FRAP analysis revealed that siRNA knockdown of 381 proteins alters nucleolar morphology and NPM1 dynamics. Consistent with this, the NPM1 dynamics-focused study also suggests that LSU precursors shape nucleolar morphology. Similarly, Quinodoz et al. proposed that LSU precursors are required to form the outermost layer of the nucleolus. How LSU precursors influence the RNA and protein composition of the nucleolus remains unclear, warranting future studies. Proximity-labeling approaches, such as those used in this current study to investigate the impact of 5’ETS retention, may provide valuable insights. Together, these findings will enhance our understanding of the biological implications of specific rRNA-processing steps on nucleolar organization and function.

## DATA AVAILABILITY

All data are available from the corresponding author upon reasonable request. All the primers and probes used in the paper are listed in Table S1. APEX2-MS proteomics data have been deposited with MassIVE MSV000098278. Any data supporting this study’s findings and additional information required to reanalyze the data reported in this paper are available in the Statistics Source Data file and from the corresponding authors upon request.

## ACKNOWLEDGEMENTS

This work was supported by the National Institutes of Health (R35 GM133721, R01 HL160726, RSG-22-064-01-RMC American Cancer Research Scholar, and Damon Runyon Innovator Award to KF. Liu. We thank Dr. Matthew Kayser for sharing the Leica SP8 confocal microscope.

## AUTHOR CONTRIBUTION

M.S.M.F. and K.F.L. designed the experiments. M.S.M.F. performed the labeling of nascent RNA, and the APEX2-Nucleolin proteomics experiments. M.S.M.F. and H.Y.T. performed proteomics analysis. M.S.M.F., C.A., and E.L. performed the FRAP, immunofluorescence imaging, and western blot experiments. M.S.M.F., A.Y.C., and H.E. generated the constructs used in this study. M.S.M.F. and Y.G. performed northern blot experiments. M.S.M.F., V.R.P., D.C., and K.F.L. wrote the manuscript. All authors participated in the discussion and editing of the manuscript.

## SUPPLEMENTAL INFORMATION

Document S1. Extended Data Figures 1-4 and Tables S1.

Table S2. NPM1-Apex2 MS Excel file.

---

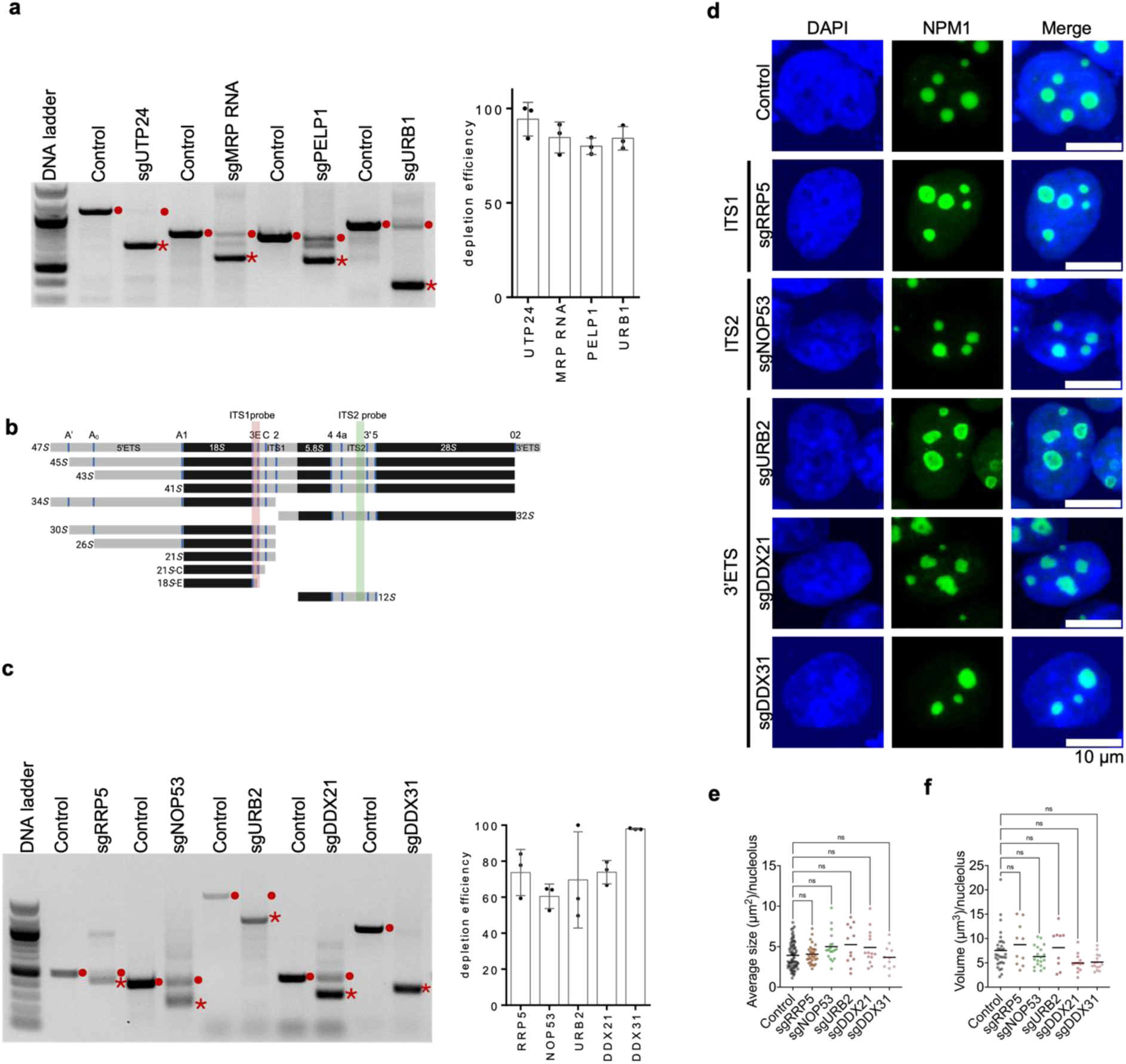

