## Supplemental figures for "Defining the impact of rRNA processing on nucleolar organization and function"

**Extended Data Figure 1. Specific pre-rRNA processing factors are critical for nucleolar morphology.**

(**a**) Depletion efficiency of ribosome assembly factors (UTP24, MRP RNA, PELP1, and URB1) in HEK293T cells was assessed using genomic PCR following CRISPR-Cas9 targeting with paired sgRNAs. Cells transfected with the empty pX459 plasmid were used as Control. PCR products from uncleaved genomic DNA are indicated by a red dot; products from cleaved DNA are marked with a red star. Right: Quantification of genomic PCR products from three biological replicates. Data represent mean ± standard deviation.

(**b**) Schematic of human pre-rRNA processing intermediates and the binding sites of DNA probes used in Northern blotting (Fig. 1c). Probe sites targeting ITS1 and ITS2 are indicated by vertical red and green lines, respectively.

(**c**) Representative gel showing depletion efficiency of additional pre-rRNA cleavage factors (targeting ITS1, ITS2, or 3′ETS regions) using CRISPR-Cas9 with paired sgRNAs. Uncleaved and cleaved genomic PCR products are marked with a red dot and red star, respectively. Right: Quantification of genomic PCR products from three biological replicates. Data represent mean ± standard deviation.

(**d**) Representative immunofluorescence images showing DAPI and Nucleophosmin (NPM1; GC marker) staining in Control and HEK293T cells depleted of ribosome assembly factors involved in cleavage of ITS1, ITS2, or 3′ETS regions. Scale bar = 10 μm.

(**e**) Quantification of nucleolar area ($\mu m^2$) from panel (d). Kruskal–Wallis test; ns = not significant ($\alpha > 0.05$). Mean values are shown.

(**f**) Quantification of nucleolar volume ($\mu m^3$) from panel (d). Kruskal–Wallis test; ns = not significant ($\alpha > 0.05$). Mean values are shown.

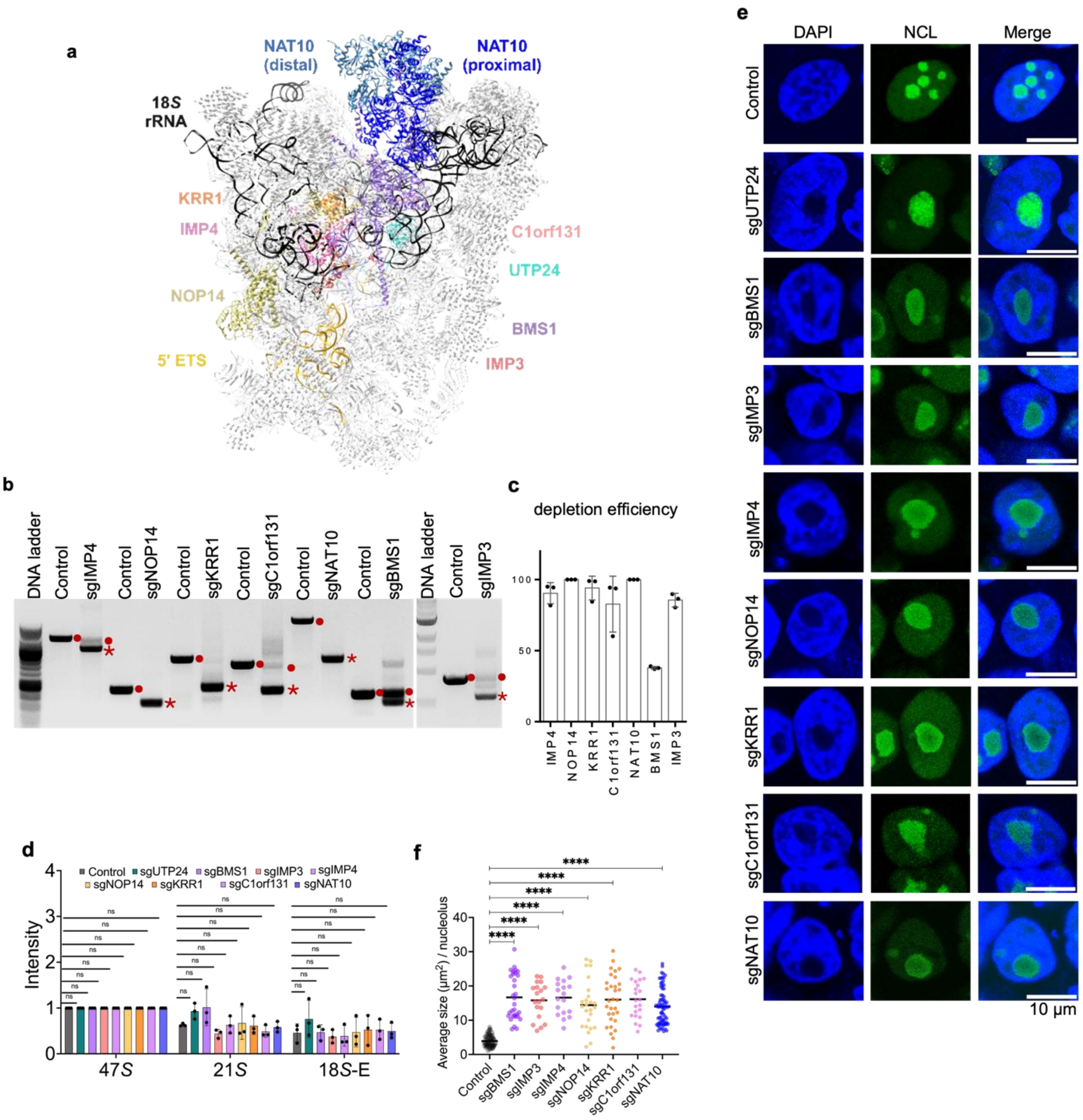

**Extended Data Figure 2. Pre-rRNA processing factors involved in A1 site cleavage are essential for nucleolar organization.**

(**a**) Structure of the human SSU processome in the pre-A1 state (PDB: 7MQ8), with protein factors implicated in A1 site cleavage highlighted in color.

(**b**) Assessment of CRISPR-Cas9–mediated depletion of A1 site cleavage factors by genomic PCR. A representative agarose gel shows depletion efficiency. The product from uncleaved genomic DNA is indicated by a red dot; the product from cleaved genomic DNA is marked with a red star.

(**c**) Quantification of PCR products shown in panel (b) across three independent biological replicates. Data represent mean ± standard deviation.

(**d**) Quantification of rRNA processing intermediates (47*S*, 21*S*, and 18*S*-E) from Fig. 2a following depletion of the indicated ribosome assembly factors. Band intensities were quantified using Fiji and normalized to the corresponding 47*S* precursor. Statistical analysis was performed using two-way ANOVA; ns = not significant ($\alpha > 0.05$). Data represent mean ± standard deviation.

(**e**) Representative immunofluorescence images of HEK293T cells transfected with either the pX459 control vector or with paired sgRNAs targeting specific ribosomal processing factors. Cells were stained with DAPI and an antibody against Nucleolin (NCL), a marker of the granular component. Scale bar: 10 µm.

(**f**) Quantification of nucleolar area ($\mu m^2$) from panel (e). Statistical analysis was performed using one-way ANOVA; ****$p < 0.0001$; ns = not significant ($\alpha > 0.05$). The data represent mean values.

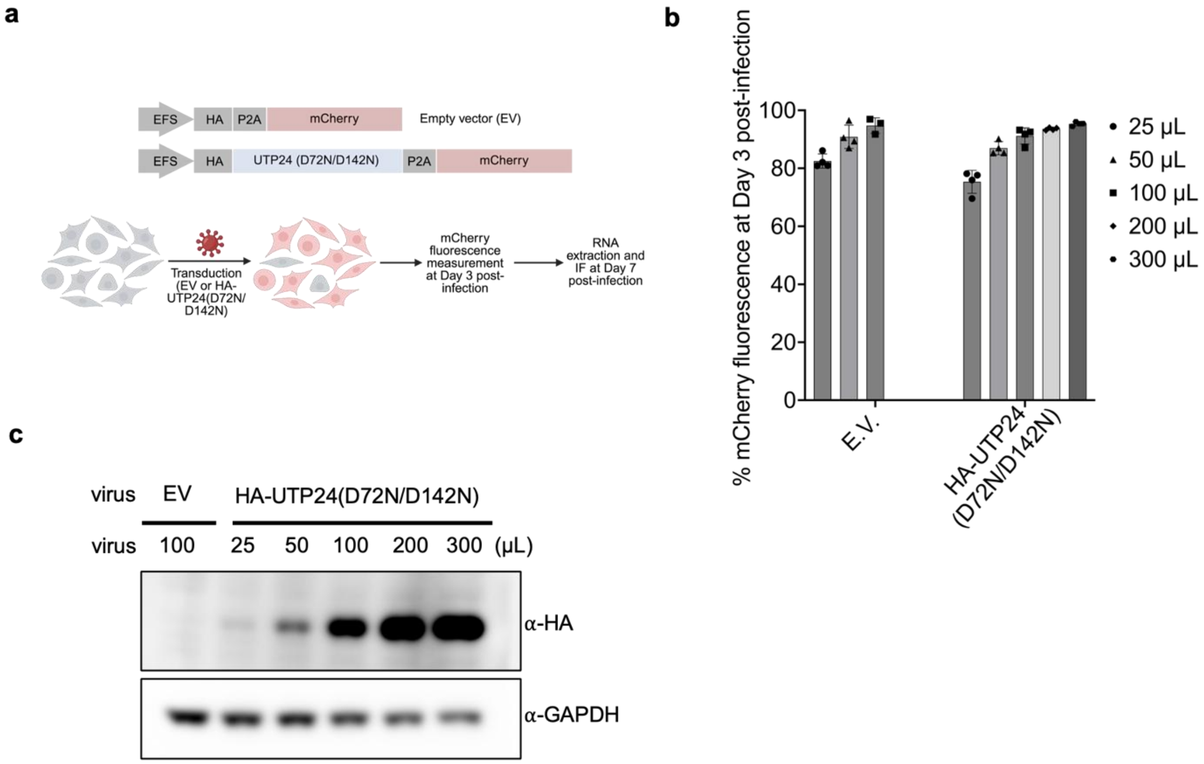


**Extended Data Figure 3. Lentiviral expression of catalytically inactive HA-UTP24(D72N/D142N).**

(**a**) Schematic of lentiviral vectors used to overexpress catalytically inactive HA-tagged UTP24(D72N/D142N). Constructs contain a P2A-mCherry cassette for monitoring transduction efficiency.

(**b**) HEK293T cells were transduced with either Lenti-P2A-mCherry (empty vector, EV) or Lenti-HA-UTP24(D72N/D142N)-P2A-mCherry.

(**c**) Transduction efficiency was quantified by measuring mCherry fluorescence at 3 days post-infection using the indicated viral doses. Data represent mean ± standard deviation.

(**d**) Expression of HA-UTP24(D72N/D142N) was confirmed by Western blot using an anti-HA antibody. GAPDH served as a loading control.

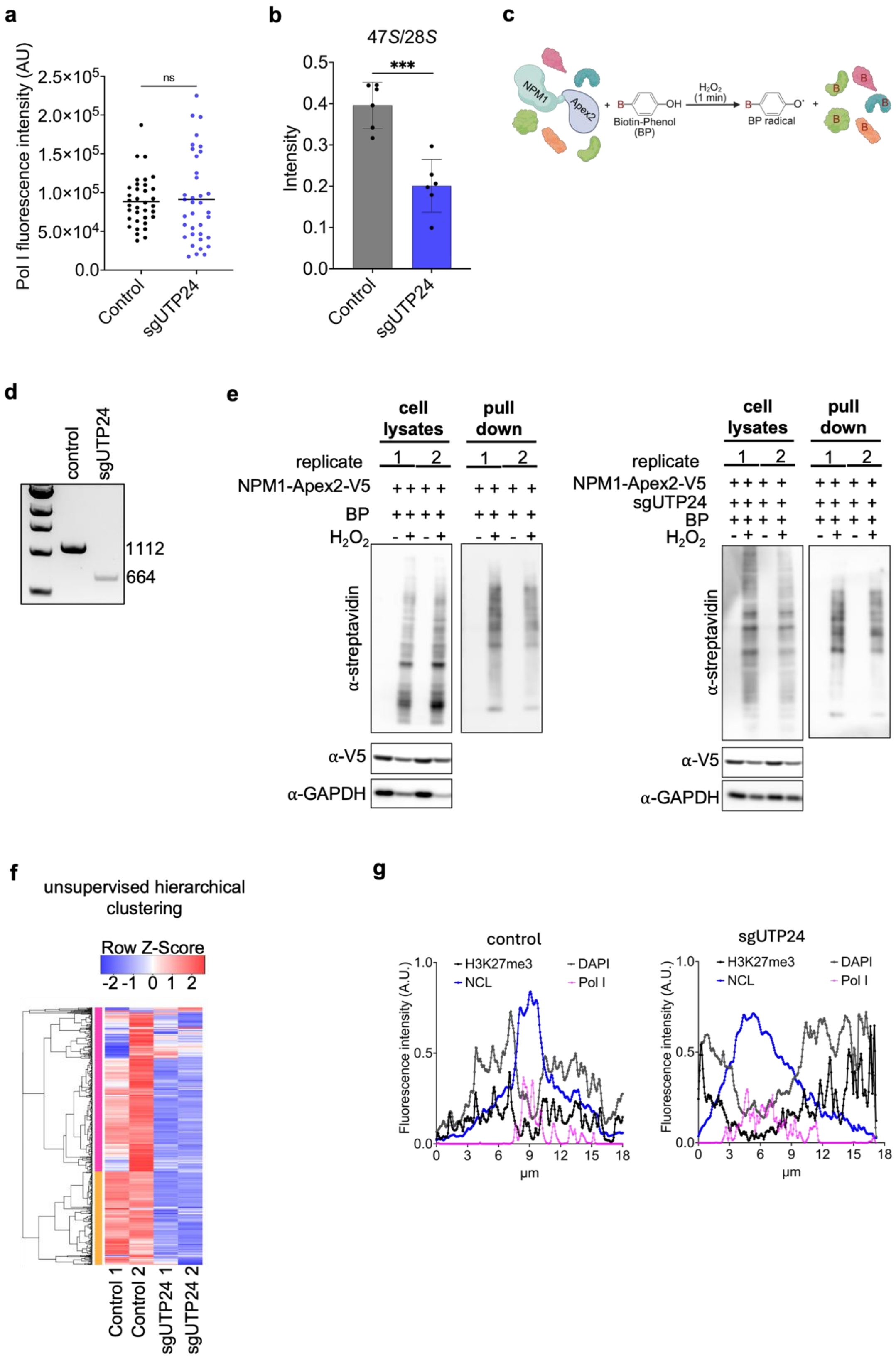

a
Pol I fluorescence intensity (AU)
2.5×10^5
2.0×10^5
1.5×10^5
1.0×10^5
5.0×10^4
0.0
ns
Control
sgUTP24
b
47S/28S
Intensity
0.5
0.4
0.3
0.2
0.1
0.0
***
Control
sgUTP24
c
NPM1
Apex2
Biotin-Phenol (BP)
H2O2 (1 min)
BP radical
d
control
sgUTP24
1112
664
e
cell lysates
pull down
replicate
1
2
NPM1-Apex2-V5
BP
H2O2
α-streptavidin
α-V5
α-GAPDH
cell lysates
pull down
replicate
NPM1-Apex2-V5
sgUTP24
BP
H2O2
α-streptavidin
α-V5
α-GAPDH
f
unsupervised hierarchical clustering
Row Z-Score
-2 -1 0 1 2
Control 1
Control 2
sgUTP24 1
sgUTP24 2
g
control
sgUTP24
H3K27me3
DAPI
NCL
Pol I
Fluorescence intensity (A.U.)
1.0
0.5
0.0
0 3 6 9 12 15 18
µm

**Extended Data Figure 4. UTP24 depletion disrupts nucleolar protein interactome and heterochromatin organization.**

(**a**) Quantification of total Pol I fluorescence intensity in morphologically normal nucleoli of control cells versus the enlarged nucleoli observed in UTP24-depleted cells. Statistical analysis was performed using an unpaired t-test; ns = not significant ($\alpha > 0.05$). Mean values are shown. AU, arbitrary units.

(**b**) Quantification of 47*S*/28*S* rRNA ratios from Northern blots shown in Figures 1c and 2a. Analysis was performed across six biologically independent replicates using Fiji software. Unpaired t-test; ***$p < 0.05$ ($\alpha > 0.05$). Mean ± standard deviation is shown.

(**c**) Schematic of Nucleophosmin (NPM1)-Apex2 proximity labeling used to biotinylate proteins in living cells.

(**d**) Genomic PCR of HEK293T cells transfected with either with either the pX459 control vector or with paired sgRNAs targeting UTP24, used to evaluate UTP24 depletion efficiency. The uncleaved PCR product is 1112 bp; the cleaved product is 664 bp.

(**e**) Western blot analysis of biotinylated proteins in total cell lysates and streptavidin-enriched pull-downs from NPM1-Apex2–expressing Control and UTP24-depleted cells.

(**f**) Unsupervised hierarchical clustering of biotinylated proteins identified by mass spectrometry from streptavidin pull-downs in Control and sgUTP24 cells. Two biological replicates were analyzed per condition.

(**g**) Quantification of fluorescence intensities for H3K27me3, NCL, Pol I, and DAPI from immunofluorescence images shown in Figure 5g.
